# Stable episodic memory and high education do not influence the rate of Alzheimer’s disease pathology as measured by plasma p-tau217

**DOI:** 10.64898/2026.04.16.718397

**Authors:** Bárbara Avelar-Pereira, Nicola Spotorno, Anna Orduña Dolado, Divya Bali, Annelie Nordin Adolfsson, Niklas Mattsson-Carlgren, Sebastian Palmqvist, Shorena Janelidze, Oskar Hansson, Lars Nyberg

## Abstract

Alzheimer’s disease (AD) neuropathological changes can be detected with blood-based biomarkers during the long preclinical phase that precedes clinical diagnosis. Tau phosphorylated at threonine 217 (p-tau217) has been found to closely correlate with brain Aβ burden. A recent large-scale cross-sectional study showed elevated p-tau217 concentrations in older individuals (Aarsland et al., 2025). This increase was higher in those with AD dementia and mild cognitive impairment (MCI), and lower in those with intact cognition and higher educational attainment. Thus, intact cognition and higher education may be associated with lower levels of AD neuropathological changes. Here we tested this hypothesis using longitudinal data from the population-based *Betula* study (n=1005; 1531 samples). The results revealed increases with increasing age over 10 years in p-tau217, where individuals with accelerated episodic-memory decline had the strongest increase. There were no differences in p-tau217 trajectories between individuals with lower or higher education or with well-maintained or age-typical decline in episodic memory. The lack of association with education was further replicated in the independent BioFINDER-2 cohort. These findings underscore the value of plasma p-tau217 for detecting early pathological changes in population-based settings but provide no support that individuals with well-maintained episodic memory or high educational attainment are spared from neuropathological changes.

## Introduction

Alzheimer’s disease (AD) is a neurodegenerative disorder characterized by the accumulation of extracellular amyloid-β (Aβ) plaques and intracellular aggregates of hyperphosphorylated tau (Jack et al., 2024). Over time, Aβ and tau pathology disrupt synaptic function and lead to neuronal loss, resulting in cognitive impairment, notably episodic-memory decline (Hardy & Selkoe, 2002; Price & Morris, 1999). AD is characterized by a long preclinical phase and there is growing interest in blood-based biomarkers that can provide diagnostic information (Hansson et al., 2023). Tau phosphorylated at threonine 217 (p-tau217) has been found to closely correlate with brain Aβ burden, discriminate AD from other neurodegenerative disorders with high accuracy, and predict cognitive decline in non-demented samples (Warmenhoven et al., 2025; Ashton et al., 2024; Palmqvist et al., 2020; Grande et al., 2025; Devanarayan et al., 2024; Mielke et al., 2022; Ashton et al., 2022).

A recent large-scale study (Aarsland et al., 2025) used 11,486 blood samples to investigate p-tau217 burden in relation to age and several other variables including cognition and education. P-tau217 concentration levels indicative of AD neuropathology were (i) higher in older age groups, (ii) higher for individuals with mild cognitive impairment (MCI) than for cognitively unimpaired individuals, and (iii) highest for older individuals with only primary education. Thus, individuals with intact cognition and higher educational attainment appear to have lower levels of p-tau217, which may reflect reduced AD pathology. Here we test this prediction using longitudinal data from the population-based *Betula* study in northern Sweden (Nyberg et al., 2020; Nilsson et al., 2004).

First, 10-year longitudinal changes in plasma p-tau217 were examined in relation to three previously defined episodic-memory aging profiles over 15 years (Josefsson et al., 2012): (i) age-typical memory decline (“average”); (ii) accelerated memory decline (“decliners”), and high-stable memory performance (“maintainers”). Based on findings that decliners have elevated risk of AD and that maintainers have reduced risk (Josefsson et al., 2023), we hypothesized that decliners would exhibit the greatest increase in p-tau217 over time and maintainers the lowest. Second, we explored whether educational attainment modified age-related longitudinal increases in plasma p-tau217. The recent study by Aarsland et al. (2025) found that higher educational attainment would be associated with a lower increase in p-tau217.

## Materials and Methods

### Participants

Data used in preparation of this work were obtained from the Betula project on aging, memory and dementia (see http://www.umu.se/en/betula), a longitudinal population-based study conducted in Umeå, Sweden (Nyberg et al., 2020). Participants were randomly sampled from the population registry and stratified by age and sex to ensure proportional representation of the general population. Exclusion criteria included dementia at baseline, major neurological or psychiatric disorders, intellectual disability, severe sensory impairment, and non-native Swedish language (Nilsson et al., 2004). To date, six main data collection waves (W1-W6; 1988–2014) have been completed at approximately five-year intervals, each including a wide range of health and cognitive assessments. Blood sampling was part of the protocol at all waves, with plasma collection introduced at wave 3 (W3; 1998–2001). The present study included 1005 participants from the longitudinal Betula cohorts S1 and S3 with available plasma samples at wave 3 (W3). All participants were aged over 50 years and had a previously defined cognitive profile based on the intercept and slope of a five-item episodic memory composite score, classifying individuals into three trajectories of episodic memory performance: maintainers, average performers, and decliners (Josefsson et al., 2012). Of these 1005 participants, 526 also had follow-up plasma samples at wave 5 (W5; 2008–2010), yielding a total of 1531 samples available for plasma p-tau217 analysis.

### Cognitive assessment and classification of cognitive aging profiles

Cognition was assessed using a composite measure of episodic memory performance. The composite included five standardized tasks: (1) immediate free recall of 16 visually and orally presented sentences, (2) delayed cued recall of nouns from the sentences, (3) immediate free recall of 16 enacted sentences, (4) delayed cued recall of nouns from enacted sentences, and (5) immediate free recall of 12 orally presented nouns. Procedures remained constant across waves. Cognitive profiles were determined following established Betula procedures and have been previously described in full detail elsewhere (see Josefsson et al., 2012; Pudas et al., 2013).

### Educational attainment

Education was assessed as self-reported years of formal education, recorded with half-year precision. For analytical purposes, education was dichotomized within each age group (i.e., 50–59, 60–69, 70–79, 80–99 years) based on the median number of years of education to account for cohort differences in educational access. Participants with years of education above the age-specific median were classified as the higher education group, whereas those with years of education equal to or below the median were classified as the lower education group.

### Plasma biomarker analyses

Venous blood samples were collected at each assessment wave from non-fasting participants during daytime hours. From W3 onward, blood collected in EDTA tubes was centrifuged to separate plasma, which was then aliquoted into secondary tubes, and stored at -80 °C pending biochemical analysis. A small subset of W3 samples (N = 94) had undergone prior thawing for earlier investigations.

Plasma p-tau217 concentrations were quantified using a validated immunoassay platform with high analytical sensitivity and specificity for AD-related phosphorylation at threonine 217 (Lilly p-tau217 in-house immunoassay; Palmqvist et al., 2020; Bali et al., 2024). The randomization procedure was based on the characteristics of the 1005 W3 participants. W3 samples were randomized according to age, sex, and cognitive profile to ensure an even distribution of participant characteristics across the 42 assay plates. For the longitudinal subset, the 526 W5 samples were positioned on the same plate and adjacent to the corresponding baseline (W3) sample to enhance within-individual robustness and reduce plate-to-plate effects. Before finalizing the plate layout, we verified that the positioning of samples did not result in an excessively skewed distribution of dementia status or cohort affiliation (S1/S3). Plasma p-tau217 concentrations were log-transformed prior to statistical analyses. Analyses were performed blinded to cognitive status.

### Statistical analyses

All statistical analyses were conducted in R (v4.4.3). To compare participants with longitudinal data to those who dropped out after W3, we used Welch’s and chi-square tests. We then fit a linear regression model (lm) with p-tau217 as the outcome variable and predictors including mean-centered age, sex, and cognitive aging profile (maintainers, average performers, and decliners). A control analysis including follow-up status (longitudinal vs. drop-out) was conducted and yielded results consistent with the main findings. We also carried out a sensitivity analysis in the subset of participants with only baseline data (dropouts) following the same procedures (a detailed description of the sensitivity analysis is provided in the Supplementary Materials).

Longitudinal changes in p-tau217 were assessed using linear-mixed effects models (lme4 package) with random intercepts for participants and fixed effects of time, mean-centered age, sex, cognitive profile, and interactions between time x age as well as time x cognitive profile. Model fit was evaluated against a simpler model excluding the time x cognitive profile interaction using likelihood ratio tests under maximum likelihood estimation. All analyses were repeated after excluding participants who developed dementia (Supplementary Materials), to assess the extent to which longitudinal p-tau217 changes were driven by individuals on a dementia trajectory. Post hoc contrasts of estimated marginal means were conducted to examine changes in p-tau217 from W3 to W5 within each cognitive profile. This was done using the emmeans package in R, with Bonferroni adjustment for multiple comparisons.

To examine whether educational attainment modified baseline levels and longitudinal changes in plasma p-tau217, an additional mixed-effects model included education as a predictor together with time, age, sex, and an interaction between time x age. Statistical significance was set at p < 0.05 (two-tailed) for all analyses. Longitudinal trajectories of plasma p-tau217 between W3 and W5 were visualized using individual participant line plots stratified by group. Group-level trends were summarized by plotting mean p-tau217 values at each time point, with shaded ribbons representing bootstrapped 95% confidence intervals. Finally, to visualize trajectories of plasma p-tau217 in relation to educational attainment, locally estimated scatterplot smoothing (LOESS) curves were overlaid for each education group. Individual data points and within-participant trajectories were retained to illustrate the distribution of observed values.

### Replication cohort: BioFINDER-2

We tested the potential association between p-tau217 levels and education in a replication cohort from the Swedish BioFINDER-2 study (https://biofinder.se/two/; Palmqvist et al., 2020). This analysis included 388 cognitively unimpaired individuals aged over 50 years who had available p-tau217 levels quantified using the same assay (see Supplementary materials for sample size details). The classification of being cognitively unimpaired has been described previously (Palmqvist et al., 2023).

## Results

### P-tau217 in the different memory-trajectory groups

In the full baseline sample (N = 1005), a linear regression analysis adjusting for age, sex, and cognitive profile showed that p-tau217 was higher in older age (β = 0.0066, SE = 0.0010, t = 6.49, p < .001). There was no significant difference between males and females (β = -0.037, SE = 0.020, t = -1.85, p = 0.065). Compared to average performers, decliners showed higher p-tau217 levels (β = 0.093, SE = 0.034, t = 2.72, p = 0.007) and maintainers showed lower levels (β = -0.069, SE = 0.024, t = -2.90, p = 0.004). We carried out a control analysis including follow-up status (longitudinal vs. drop-out), with results indicating that those who remained in the study did not differ significantly at baseline from those who dropped out (β = -0.011, SE = 0.0218, t = -0.50, p = 0.618; Supplementary Materials). As a sensitivity analysis, the cross-sectional model was repeated in the drop-out sample only, yielding similar albeit attenuated results. Comparable findings were also observed after excluding participants with dementia, indicating that baseline differences were not primarily driven by dementia cases (Supplementary Materials).

The main analysis considered longitudinal increases in p-tau217 in the three memory-trajectory groups. A linear mixed-effects model (Table 1) examined changes in p-tau217 over time. The model revealed a significant effect of time, such that p-tau217 levels were higher at follow-up compared to baseline across the sample (β = 0.179, SE = 0.016, t(646.8) = 11.11, p < 0.001). Baseline age was positively associated with p-tau217 (β = 0.0066, SE = 0.0011, t(1243) = 6.25, p < 0.001), indicating that older participants exhibited higher levels. Moreover, there was a significant time x age interaction (β = 0.0119, SE = 0.0015, t(686.4) = 7.88, p < 0.001), suggesting that the increase in p-tau217 from W3 to W5 was more pronounced in older participants. Sex was not significantly associated with p-tau217 (β = -0.037, SE = 0.020, t(975.6) = -1.87, p = 0.061).

**Table 1.** Results of linear mixed-effects model for longitudinal increases in plasma p-tau217.

| Predictor | Estimate | Std. Error | df | t value | p value |
| --- | --- | --- | --- | --- | --- |
| Intercept | -1.138 | 0.0161 | 1148 | -70.72 | < 2e-16 *** |
| Time | 0.179 | 0.0161 | 646.8 | 11.11 | < 2e-16 *** |
| Decliners (vs Average) | 0.0930 | 0.0355 | 1243 | 2.62 | 0.00894 ** |
| Maintainers (vs Average) | -0.0688 | 0.0246 | 1240 | -2.80 | 0.00526 ** |
| Age | 0.00660 | 0.00106 | 1243 | 6.25 | 5.70e-10 *** |
| Sex | -0.0371 | 0.0198 | 975.6 | -1.87 | 0.0613 |
| Time x Decliners | 0.1555 | 0.0433 | 639.2 | 3.59 | 0.000357 *** |
| Time x Maintainers | -0.00417 | 0.0289 | 627.6 | -0.145 | 0.8851 |
| Time x Age | 0.0119 | 0.00151 | 686.4 | 7.88 | 1.28e-14 *** |

\*\*\*p < 0.001, \*\*p < 0.01, \*p < 0.05.

Critically, the results showed an interaction between time and cognitive profile, with decliners exhibiting significantly larger increases from W3 to W5 compared to average performers (β = 0.155, SE = 0.043, t(639.2) = 3.59, p < 0.001; Figure 1). However, average performers and maintainers did not differ significantly in their change over time (β = -0.004, SE = 0.029, t(627.6) = -0.15, p = 0.885). Overall, these results confirm that p-tau217 increases longitudinally across all cognitive profiles, with the steepest rise for decliners. Maintainers showed numerically lower values than average performers, but this difference was not statistically significant. Sensitivity analyses excluding participants who developed dementia indicated that the longitudinal differences between cognitive profiles were significantly attenuated. Specifically, decliners and average performers did not differ in their levels of p-tau217 (β = 0.0387, SE = 0.0482, t = 0.80, p = 0.423) and maintainers showed only a trend toward lower levels (β = –0.0465, SE = 0.0260, t = –1.79, p = 0.074). This suggests that the observed increase in p-tau217 among decliners likely reflects emerging dementia pathology (Supplementary Materials for full model results).

**Figure 1.**
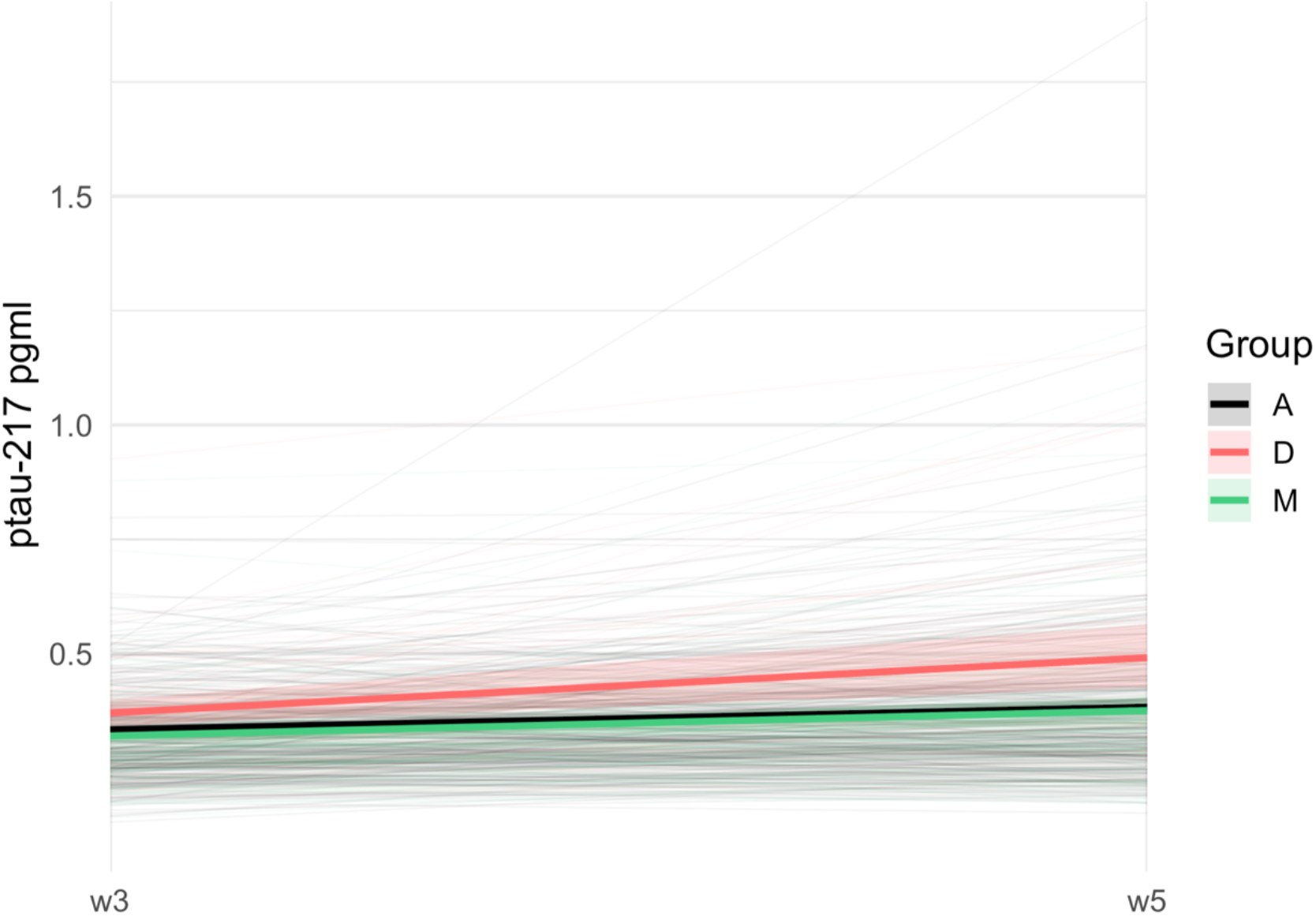
Longitudinal trajectories of p-tau217 in participants longitudinally characterized as maintainers, average performers, and decliners in episodic memory.

### P-tau217 in relation to educational attainment

Baseline differences in p-tau217 between the higher and lower educational groups were small in all age cohorts (Table 2). Also, a linear mixed-effects model examining longitudinal associations with educational attainment (Table 3) revealed no significant relationship between education and overall p-tau217 levels (β = 0.0105, SE = 0.0076, t(1425) = 1.38, p = 0.168). The longitudinal trajectories indicated substantial overlap between educational groups over time (Figure 2; see Supplementary Materials for sex-stratified plots). These findings provide no evidence for an association between educational attainment and plasma p-tau217.

**Table 2.**
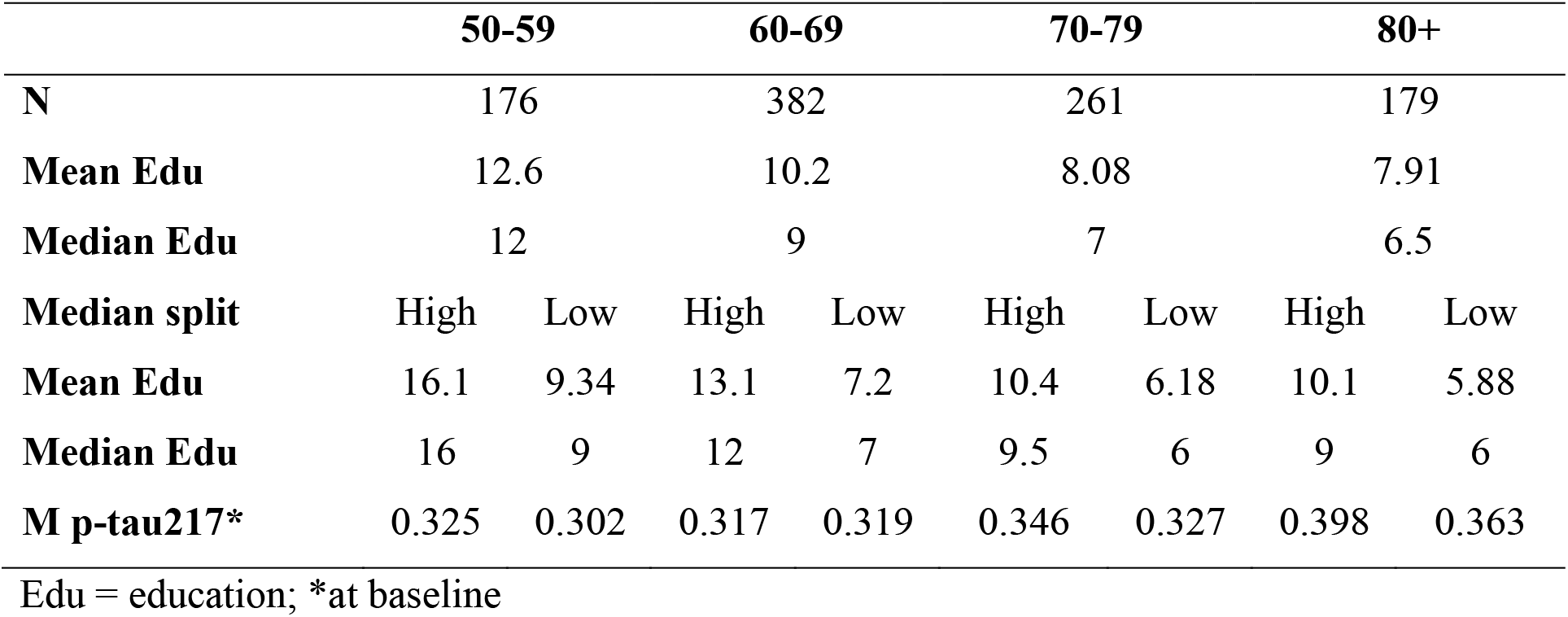
Educational attainment and baseline plasma p-tau217 across age groups.

**Table 3.** Results of linear mixed-effects model examining associations between educational attainment and longitudinal changes in plasma p-tau217.

| Predictor | Estimate | Std. Error | df | t value | p value |
| --- | --- | --- | --- | --- | --- |
| Intercept | 0.336 | 0.0163 | 1208 | 20.64 | < 2e-16 *** |
| Time | 0.0183 | 0.0070 | 1265 | 2.60 | 0.00934 ** |
| Age | 0.00240 | 0.00044 | 1319 | 5.43 | 6.8e-08 *** |
| Education group | 0.0105 | 0.0076 | 1425 | 1.38 | 0.16774 |
| Sex | -0.0090 | 0.0081 | 993.4 | -1.10 | 0.27043 |
| Time x Age | 0.00618 | 0.00068 | 761.6 | 9.10 | < 2e-16 *** |

\*\*\*p < 0.001, \*\*p < 0.01, \*p < 0.05.

**Figure 2.**
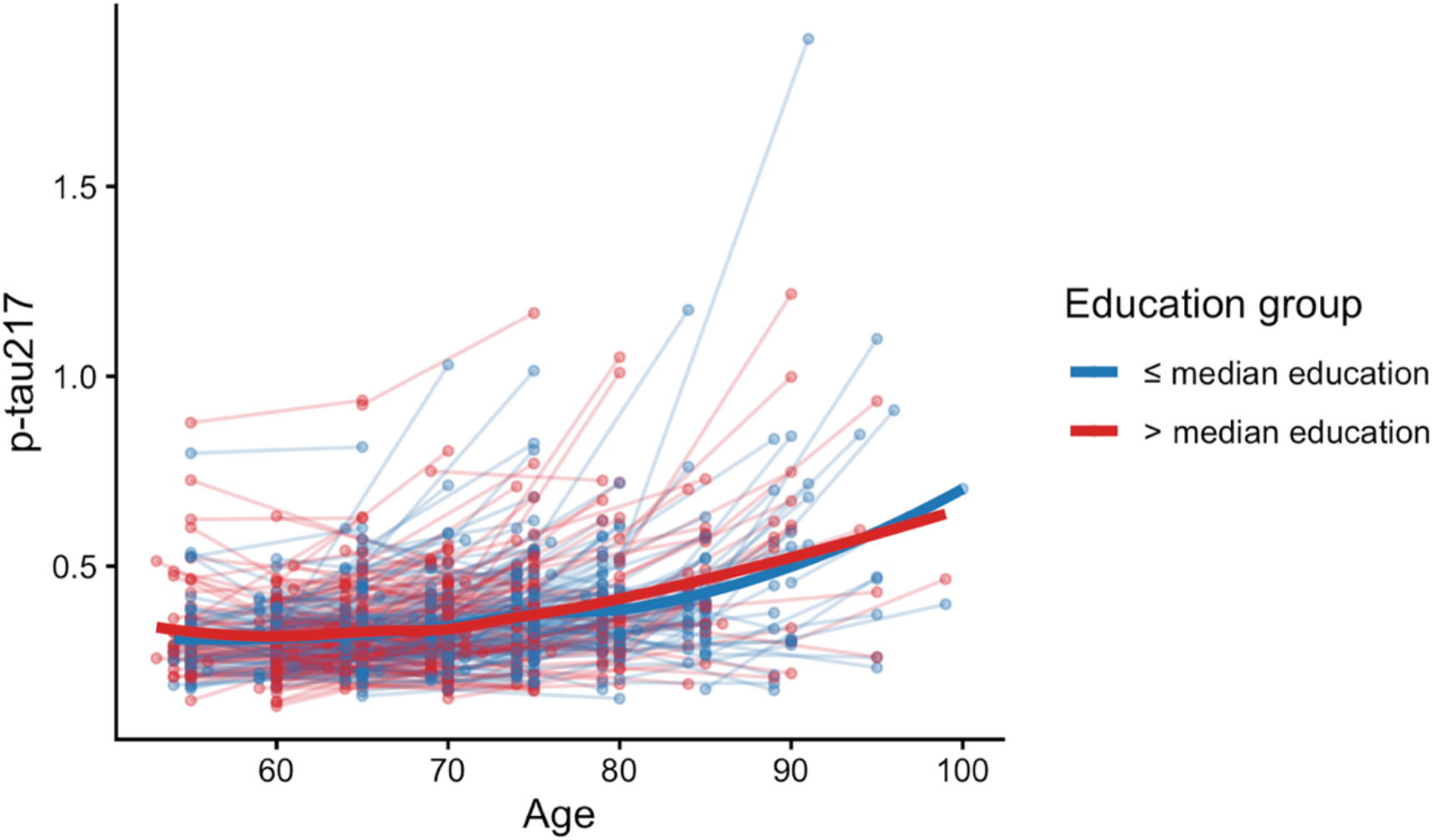
Longitudinal changes in plasma p-tau217 as a function of age. Thin lines = individuals, bold lines = mean trajectories for higher and lower education groups.

### Replication: Educational attainment and p-tau217 in BioFINDER-2

In line with the main findings, differences in p-tau217 between individuals with higher and lower education were small (Table 4). A linear regression model (Table 5) demonstrated a positive association between age and p-tau217 levels after controlling for sex (β=0.009, SE=0.002, t=4.24, p<0.001). Including an interaction term for educational level - coded as a binary variable (low vs. high) based on a median split - revealed no significant moderation effect of education on the relationship between age and p-tau217 (age x education: β=−0.006, SE=0.004, t=-1.492, p=0.14). Additionally, sex showed no statistical association with p-tau217 levels (β=−0.03, SE=0.05, t=−0.718, p>0.4). The distribution of p-tau217 across educational groups is shown in Figure 3.

**Table 4.** Educational attainment and plasma p-tau217 across age groups in the BioFINDER-2 cohort.

|  | **50-60** |  | **60-70** |  | **70-80** |  | **80+** |  |
| --- | --- | --- | --- | --- | --- | --- | --- | --- |
| **N** | 123 |  | 82 |  | 120 |  | 63 |  |
| **Mean Edu** | 13.7 |  | 12.6 |  | 11.8 |  | 12.6 |  |
| **Median Edu** | 13.0 |  | 12.0 |  | 11.0 |  | 12.5 |  |
| **Median split** | High | Low | High | Low | High | Low | High | Low |
| **Mean Edu** | 16.3 | 11.2 | 14.8 | 10.8 | 14.8 | 9.4 | 16.2 | 9.0 |
| **Median Edu** | 16.0 | 11.0 | 14.0 | 11.0 | 14.0 | 10.0 | 15.0 | 9.0 |
| **M p-tau217** | 0.162 | 0.147 | 0.180 | 0.190 | 0.203 | 0.202 | 0.197 | 0.240 |

**Table 5.**
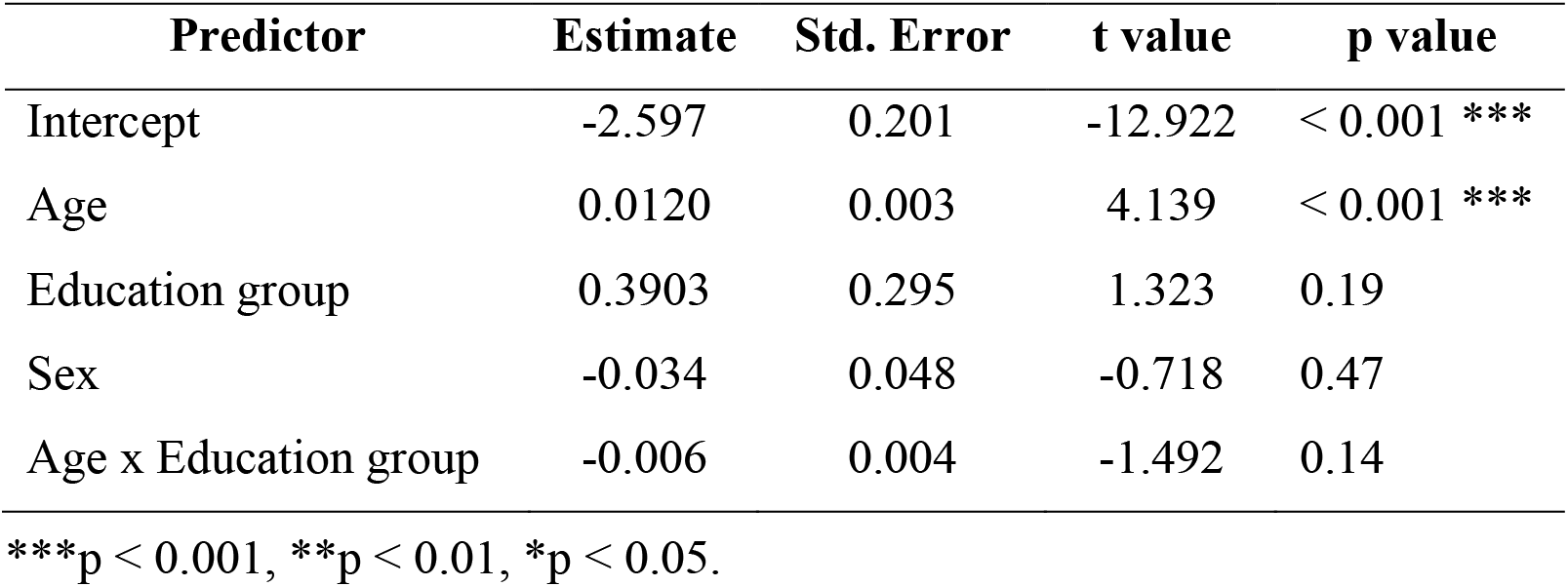
Linear regression model examining the association between educational attainment and plasma p-tau217 in the BioFINDER-2 cohort.

| **Predictor** | **Estimate** | **Std. Error** | **t value** | **p value** |
| --- | --- | --- | --- | --- |
| Intercept | -2.597 | 0.201 | -12.922 | < 0.001 *** |
| Age | 0.0120 | 0.003 | 4.139 | < 0.001 *** |
| Education group | 0.3903 | 0.295 | 1.323 | 0.19 |
| Sex | -0.034 | 0.048 | -0.718 | 0.47 |
| Age x Education group | -0.006 | 0.004 | -1.492 | 0.14 |

\*\*\*p < 0.001, \*\*p < 0.01, \*p < 0.05.

**Figure 3.**
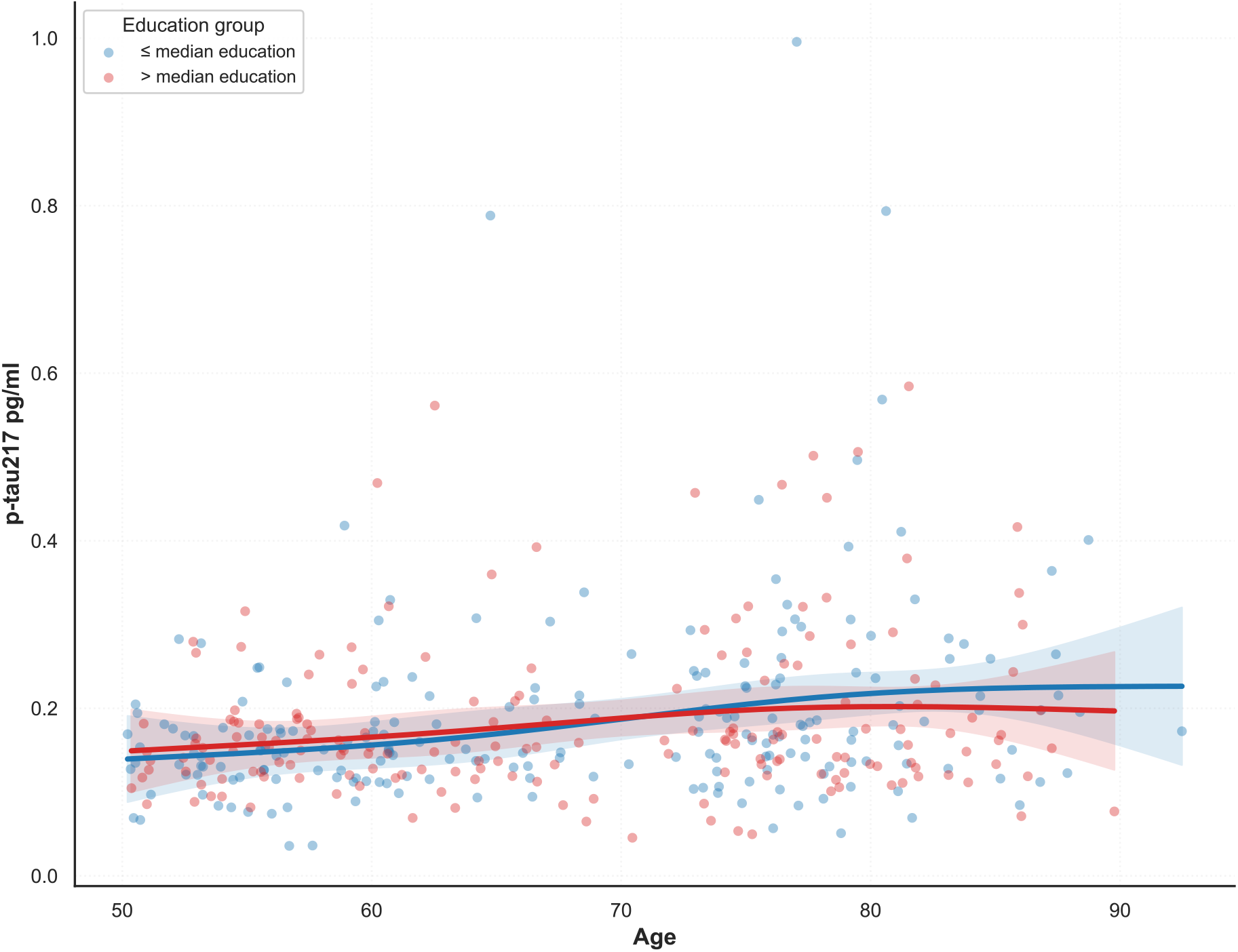
Plasma p-tau217 as a function of age in the replication cohort. Points represent individual baseline observations, and solid lines represent the fit of a natural cubic spline stratified by educational attainment. Shaded areas depict the 95% confidence intervals.

## Discussion

In this study we examined possible protective influences of well-preserved episodic memory and educational attainment on AD biomarkers. Specifically, we investigated 10-year longitudinal changes in plasma p-tau217 in relation to different longitudinal episodic-memory trajectories and variation in educational attainment.

We found that p-tau217 increased significantly over 10 years in all memory-trajectory groups, and that the increase was influenced by baseline age with a steeper increase for older participants. The p-tau217 increase was significantly greater for memory decliners compared to participants with an age-typical (average) memory trajectory. After excluding incident dementia cases, this longitudinal difference was markedly attenuated, suggesting that the accelerated p-tau217 increase among memory decliners was driven by pathology. These observations are consistent with past findings of a higher p-tau217 burden in older individuals with MCI and dementia (Aarsland et al., 2025; Milà-Alomà et al., 2022; Mattsson-Carlgren et al., 2023; Devanarayan et al., 2024; Petersen et al., 2026). Differences between memory maintainers and average performers have been demonstrated for genetic, epigenetic, and lifestyle factors (Josefsson et al., 2012; Degerman et al., 2017). Here we found that longitudinal p-tau217 trajectories overlapped for these groups, which is in line with findings of no cross-sectional differences in p-tau181 burden between Superagers and age-typical controls (Garo-Pascual et al., 2023).

For educational attainment, we found highly similar increases in plasma p-tau217 levels over the 10-year follow-up for individuals with higher or lower education. These findings were replicated in an independent BioFINDER-2 cohort of cognitively unimpaired individuals, in which no association between p-tau217 and education was also observed. As noted in the Introduction, Aarsland et al. (2025) did find that individuals with higher education had a lower prevalence of plasma p-tau217 verified AD, in particular in the oldest age groups. They noted that their findings supported a protective influence of education, for instance by contributing to a *cognitive reserve* (Stern, 2002; Stern et al., 2020). In contrast, our finding of no protective influences of education on p-tau217 build-up is consistent with those that education does not influence the rate of age-related cognitive decline (Lövdén et al., 2020; Seblova, Berggren, & Lövdén, 2020) or brain atrophy (Nyberg et al., 2021; Fjell et al., 2025), and only has modest influences on initial neurocognitive levels. These and related findings (Seblova et al., 2020) suggest that higher education supports initial brain and cognitive development but does not protect against detrimental age-related brain changes.

Several factors may help explain the discrepant findings between the present study and the work by Aarsland et al., (2025). First, their analyses focused on cross-sectional prevalence of biomarker positivity using predefined plasma p-tau217 cut-offs to classify individuals with AD. Our analyses examined continuous biomarker levels and within-person longitudinal change. Second, differences in the age composition of the samples may contribute to divergent results. In Aarsland et al. (2025), the analyses of education in relation to AD were conducted in participants aged 70 years and older. In contrast, our sample included participants from the early 50s onward. Education-related differences may therefore be more difficult to detect in younger groups where AD pathology is less common and only become apparent at older ages when pathological burden increases. Moreover, neither study fully accounted for potential confounding influences such as comorbid medical conditions, which may vary between populations and influence p-tau217 levels or their relationship with education.

The Betula project provides a unique combination of population-based data, comprehensive cognitive assessments and repeated blood sampling across multiple waves, enabling the examination of longitudinal cognitive and biomarker trajectories (Nyberg et al., 2020). Still, our study comes with certain limitations. Biomarker analyses were limited to two timepoints approximately a decade apart, restricting assessment of nonlinear changes. Future studies should combine plasma p-tau217 with neuroimaging measures such as hippocampal atrophy and indices of vascular–metabolic burden. In addition, examining how p-tau217 interacts with APOE genotype and lifestyle factors will be important.

In summary, our findings underscore the value of plasma p-tau217 for detecting early pathological changes in population-based settings but provide no support that individuals with well-maintained episodic memory or high educational attainment are protected against p-tau217 accumulation.

## Acknowledgments

We thank the Betula group and study participants, and Betula coordinator Mikael Stiernstedt for expert assistance.

## Competing Interests

OH is an employee of Lilly and Lund University. SP has acquired research support (for the institution) from Avid Radiopharmaceuticals and ki elements through ADDF. In the past 3 years, he has received consultancy/speaker fees from BioArtic, Danaher, Eisai, Eli Lilly, Novo Nordisk, Roche, and Sanofi.

## Data availability

Due to legislation and ethical restrictions, the data are not publicly available. Access may be granted upon reasonable request to the corresponding author, approval by the Betula Steering Group (umu.se/en/betula) and a confidentiality assessment.

## Funding

Supported by a scholar-grant to L.N. from the Knut and Alice Wallenberg (KAW) Foundation. This work was partly supported by a grant from the Swedish Research Council to B. A.-P. (Grant No.: 2025-00680). The BioFINDER study was funded by the National Institute of Aging (grant no. R01AG083740), European Research Council (grant no. ADG-101096455), Alzheimer’s Association (grant nos. ZEN24-1069572 and SG-23-1061717), GHR Foundation, Michael J. Fox Foundation (grant no. MJFF-025507), Lilly Research Award Program, WASP and DDLS Joint call for research projects (grant no. WASP/DDLS22-066), Swedish Research Council (grant nos. 2021-02219, 2022-00775 and 2018-02052), ERA PerMed (grant no. ERAPERMED2021-184), Knut and Alice Wallenberg Foundation (grant no. 2022-0231), Strategic Research Area MultiPark (Multidisciplinary Research in Parkinson’s disease) at Lund University, Swedish Alzheimer Foundation (grant nos. AF-1011949, AF-994229 and AF-980907), Swedish Brain Foundation (grant nos. FO2023-0163, FO2024-0284 and FO2021-0293), Parkinson foundation of Sweden (grant no. 1412/22), Cure Alzheimer’s fund, Rönström Family Foundation (grant nos. FRS-0003, FRS-0004, FRS-0011 and FRS-0013), Berg Family Foundation, Ingvar Kamprad Foundation (grant no. 20243058), Avid Pharmaceuticals, Bundy Academy, Konung Gustaf V:s och Drottning Victorias Frimurarestiftelse, Skåne University Hospital Foundation (grant no. 2020-O000028), Regionalt Forskningsstöd (grant nos. 2022-1259 and 2022-1346) and Swedish federal government under the ALF agreement (2022-Projekt0080, 2022-Projekt0107).

## Notes

### Competing Interest Statement

Oskar Hansson (OH) is an employee of Lilly and Lund University. Sebastian Palmqvist (SP) has acquired research support (for the institution) from Avid Radiopharmaceuticals and ki elements through ADDF. In the past 3 years, he has received consultancy/speaker fees from BioArtic, Danaher, Eisai, Eli Lilly, Novo Nordisk, Roche, and Sanofi.

