## Supplementary for "Stable episodic memory and high education do not influence the rate of Alzheimer’s disease pathology as measured by plasma p-tau217"

**Supplementary Materials**

**Baseline differences in p-tau217 between drop-out and longitudinal participants**

Our study includes a total of 1531 plasma samples from 1005 participants across baseline (W3; N = 1005) and follow-up (W5; N = 526). As shown in Supplementary Table 1, participants with longitudinal data were younger (Welch's $t(921.09) = 13.12$, $p < 0.001$, 95% CI [6.51, 8.80]) and showed higher baseline p-tau217 levels compared to those who dropped out after baseline (Welch's $t(935.16) = 2.98$, $p = .003$, 95% CI [0.021, 0.102]). Sex distribution did not differ between the two groups ($\chi^2(1) = 1.30$, $p = 0.25$).

Post-hoc Dunn tests (adjusted for multiple comparisons) comparing baseline age across cognitive aging profiles showed that maintainers were significantly older than average performers ($Z = -3.82$, $p < 0.001$) and decliners ($Z = -2.24$, $p = 0.0377$). However, age did not differ significantly between decliners and average performers ($Z = -0.16$, $p = 0.874$).

Supplementary Table 1. Baseline demographics and biomarker characteristics.

| **Drop-out sample** | **Maintainers** | **Average** | **Decliners** |
|---|---|---|---|
| N | 114 | 315 | 50 |
| P-tau217 ± SD | 0.342 ± 0.155 | 0.344 ± 0.143 | 0.390 ± 0.164 |
| Age (years, mean ± SD) | 74.482 ± 9.677 | 71.498 ± 10.148 | 72.420 ± 10.182 |
| Range | 54-90 | 54-91 | 54-90 |
| Sex (f/m) | 68/46 | 159/156 | 29/21 |
| **Longitudinal sample** | | | |
| N | 142 | 335 | 49 |
| P-tau217 ± SD | 0.302* ± 0.096 | 0.323 ± 0.101 | 0.351 ± 0.123 |
| Age (years, mean ± SD) | 66.965** ± 8.747 | 63.925** ± 8.033 | 62.918** ± 6.075 |
| Range | 54-90 | 53-90 | 53-80 |
| Sex (f/m) | 99/43 | 177/158 | 25/24 |

***p < 0.001, **p < 0.01, *p < 0.05. Group differences refer to comparisons between participants with longitudinal follow-up and those who contributed only baseline data (drop-outs), conducted separately within each cognitive aging profile.

Drop-outs showed higher baseline p-tau217 levels (M = 0.348 vs. 0.320) compared to participants with longitudinal data. This difference was statistically significant in an unadjusted comparison (Welch's $t(935.16) = 2.98$, $p = .003$, 95% CI [0.021, 0.102]). Interestingly, this pattern was driven by maintainers (Welch's $t(199.61) = 2.01$, $p = .046$, 95% CI [0.002, 0.170]) but not average performers (Welch's $t(628.90) = 1.84$, $p = .066$, 95% CI [–0.003, 0.095]) or decliners (Welch's $t(92.51) = 1.09$, $p = .279$, 95% CI [–0.063, 0.215]), for whom p-tau217 levels did not differ significantly between drop-outs and longitudinal participants.

When age, sex, cognitive profile, and follow-up status (drop-out vs. longitudinal) were entered into a linear regression predicting baseline p-tau217, follow-up status was not significant. Age was still a significant positive predictor ($\beta = 0.0064$, SE = 0.0011, $t = 5.76$, $p < 0.001$), indicating that older participants had higher p-tau217 levels. Compared to average performers, decliners had higher p-tau217 ($\beta = 0.0928$, SE = 0.0342, $t = 2.71$, $p = 0.0068$), whereas maintainers had lower p-tau217 ($\beta = -0.0679$, SE = 0.0238, $t = -2.85$, $p = 0.0044$). Sex was not a significant predictor ($\beta = -0.0380$, SE = 0.0203, $t = -1.87$, $p = 0.0619$). Importantly, participants who remained in the study did not differ significantly from those who dropped out at baseline ($\beta = -0.0109$, SE = 0.0218, $t = -0.50$, $p = 0.618$). Taken together, these results suggest that baseline differences in p-tau217 were not driven by follow-up status.

**Sensitivity analysis: Linear regression accounting for baseline differences in p-tau217 in the drop-out sample only**

We repeated the cross-sectional linear regression model, including age, sex, and cognitive profile, using only the drop-out sample. Age remained a significant predictor ($\beta = 0.0076$, SE = 0.0016, $t = 4.83$, $p < 0.001$), but group differences were attenuated in this smaller subsample: decliners tended to have higher p-tau217 than average performers, but this did not reach significance ($\beta = 0.0974$, SE = 0.0524, $t = 1.86$, $p = 0.064$), and maintainers did not differ significantly from average performers ($\beta = -0.0501$, SE = 0.0379, $t = -1.32$, $p = 0.187$). Sex was not associated with p-tau217 ($\beta = -0.0610$, SE = 0.0316, $t = -1.93$, $p = 0.054$). These differences likely reflect reduced statistical power.

**Exclusion of dementia cases**

We repeated all analyses after excluding participants with a dementia diagnosis. Participants with longitudinal plasma data remained significantly younger than those who contributed only baseline samples (Welch's t-test, $p < 0.001$), indicating selective attrition unrelated to dementia status. Drop-outs also showed slightly higher baseline plasma p-tau217 levels than longitudinal participants ($p = 0.032$).

Consistent with the main analyses, post-hoc Dunn tests showed that maintainers were significantly older than both average performers and decliners ($p < 0.001$ for both comparisons), whereas average performers and decliners did not differ significantly in age ($p = 0.39$).

In cross-sectional linear regression models adjusted for age, sex, and cognitive profile, plasma p-tau217 increased significantly with age ($\beta = 0.00257$, SE = 0.00040, $t = 6.44$, $p < 0.001$) but did not differ by sex ($\beta = –0.0112$, SE = 0.00794, $t = –1.41$, $p = 0.159$). Relative to average performers, decliners exhibited higher baseline p-tau217 concentrations (($\beta = 0.0364$, SE = 0.0134, $t = 2.72$, $p = 0.0067$), whereas maintainers showed lower levels ($\beta = –0.0217$, SE = 0.00928, $t = –2.34$, $p = 0.0195$).

However, the longitudinal mixed-effects model indicated that differences between cognitive profiles were attenuated after exclusion of dementia cases: decliners showed numerically higher levels than average performers ($\beta = 0.0387$, SE = 0.0482, $t = 0.80$, $p = 0.423$), but this effect did not reach statistical significance, and maintainers showed a trend toward lower levels ($\beta = –0.0465$, SE = 0.0260, $t = –1.79$, $p = 0.074$). Plasma p-tau217 increased significantly from baseline to follow-up ($\beta = 0.136$, SE = 0.0169, $t = 8.05$, $p < 0.001$), with older age associated with higher overall levels ($\beta = 0.00461$, SE = 0.00115, $t = 4.00$, $p < 0.001$) and a stronger longitudinal increase (time x age interaction: $\beta = 0.00917$, SE = 0.00164, $t = 5.59$, $p < 0.001$). Sex was not significantly associated with p-tau217 ($\beta = –0.00618$, SE = 0.0216, $t = –0.29$, $p = 0.775$).

**Longitudinal changes in p-tau217: model comparison**

We tested a simplified model without the interaction between time and cognitive profile. Results were consistent across the two models, but model fit indices indicated that the more complex model provided better fit ($\chi^2(2) = 13.51$, $p = 0.001$, $p < 0.001$: AIC = -733.70 vs. -743.21).

**BioFINDER-2 cohort**

Supplementary Table 2. Baseline characteristics of the BioFINDER-2 cohort

| **Variable** | **BioFINDER-2 (N=388)** |
|---|---|
| Age, years | 68 (11) |
| Age, range | 50 - 92 |
| Sex, n. females (%) | 219 (56%) |
| Education, years | 13 (4) |
| p-tau217 levels (pg/ml, std.) | 0.186 (0.108) |

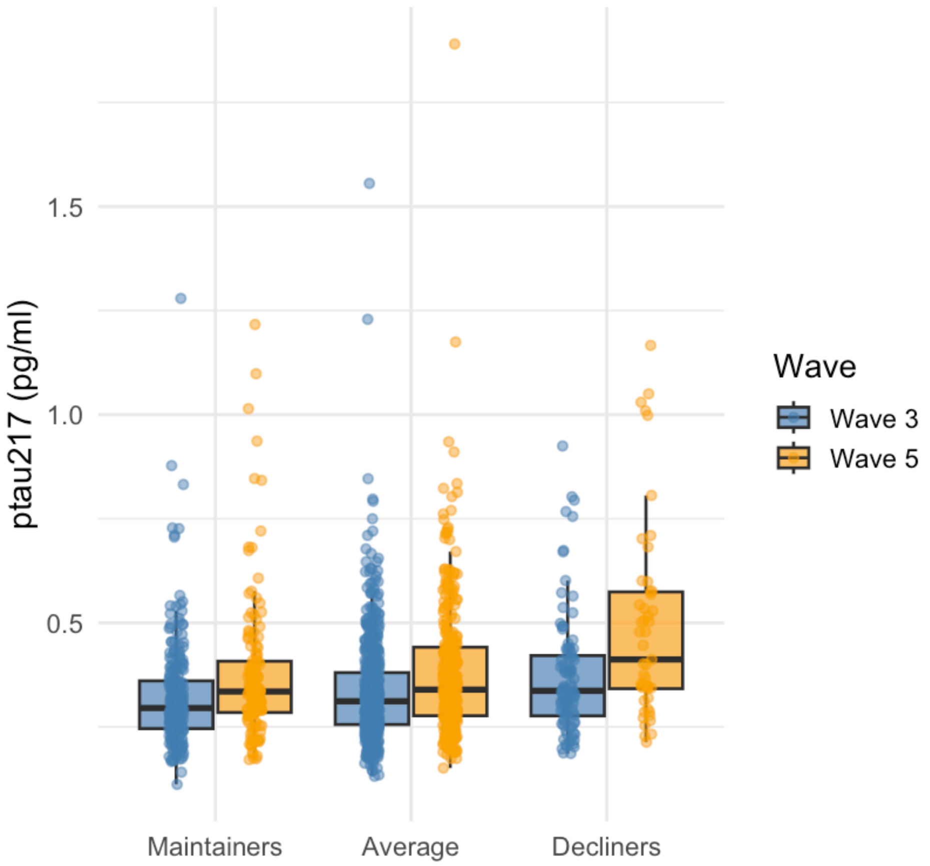


**Supplementary Figure 1.** Baseline and follow-up differences in p-tau217 among maintainers, average performers, and decliners.

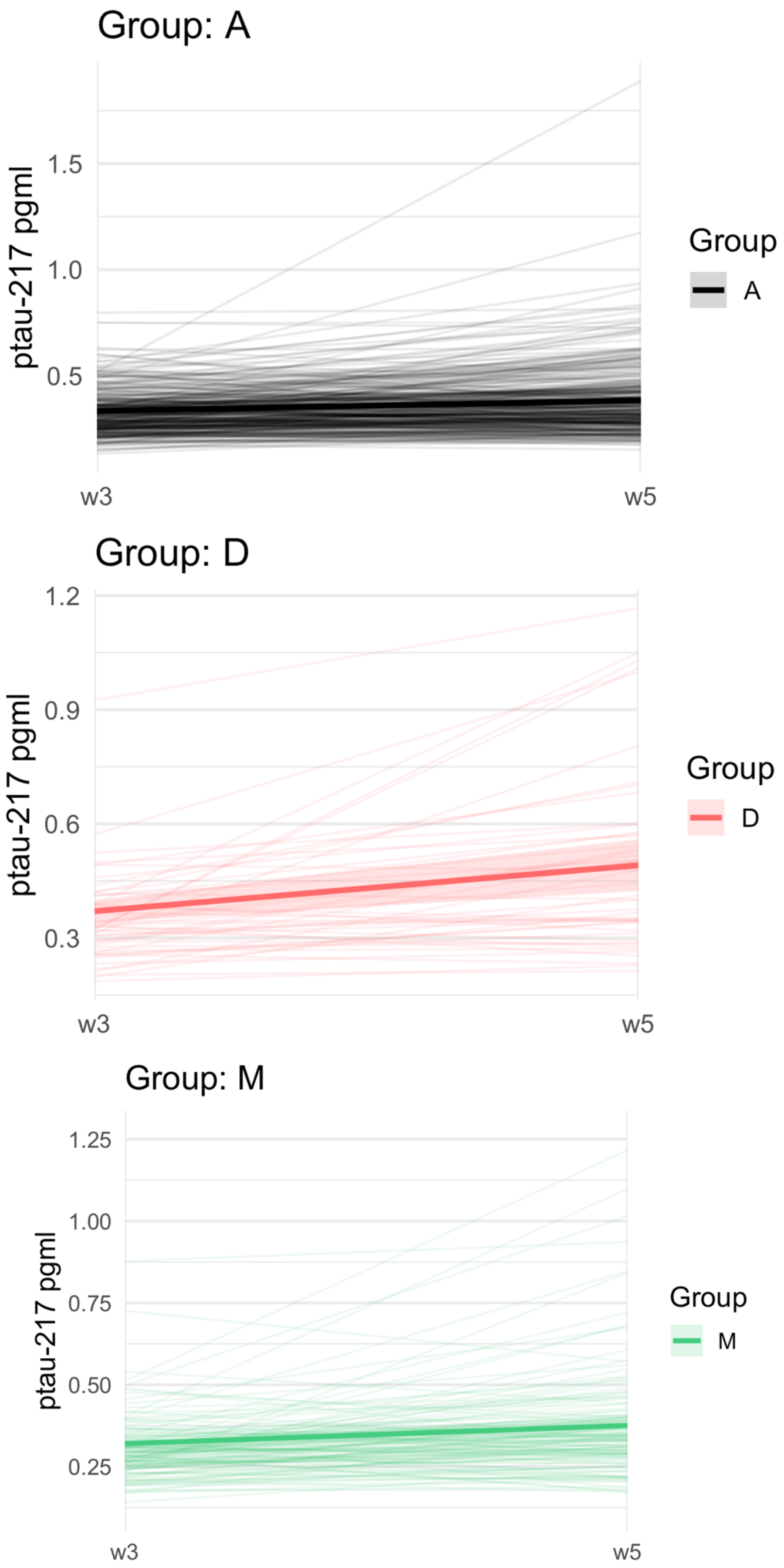


**Supplementary Figure 2.** Trajectories of p-tau217 for each cognitive profile.

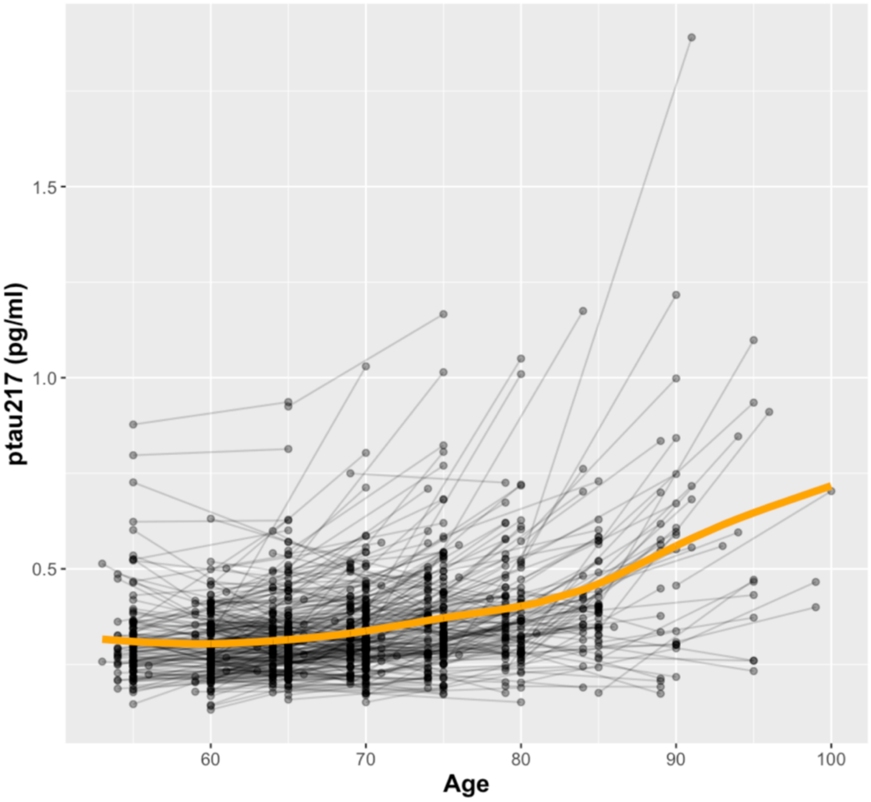


**Supplementary Figure 3.** Individual trajectories for p-tau217 changes. The bold orange line indicates mean change, estimated using a Generalized Additive Mixed Model.

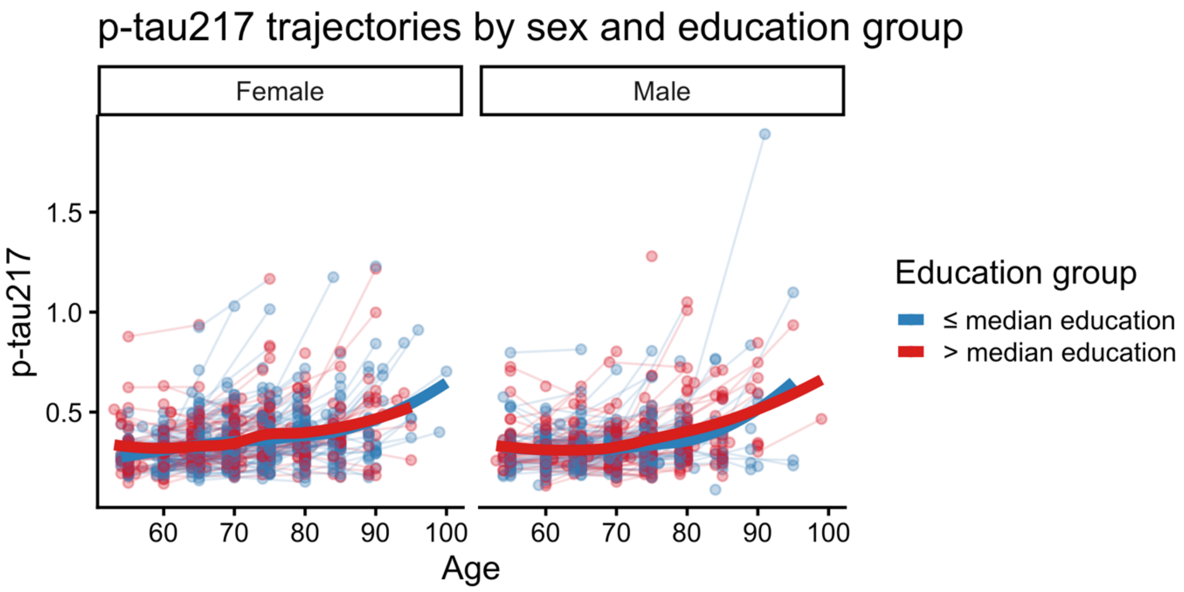


**Supplementary Figure 4.** Individual and mean trajectories of plasma p-tau217 by sex.
